# Analyte-Class-Dependent Electrophoretic Organization Enables Single-Run Proteome–Metabolome Analysis by CE–ESI–MS

**DOI:** 10.64898/2026.07.03.736334

**Authors:** Ramesh Kumar, Ryan O’Neal, Peter Nemes

**Affiliations:** Department of Chemistry & Biochemistry and; Brain and Behavior Institute, University of Maryland, College Park, MD 20742, United States

**Keywords:** capillary electrophoresis, mass spectrometry, metabolomics, proteomics, single cell, *Xenopus laevis*

## Abstract

Dual proteome–metabolome measurements from limited samples typically require sample splitting or sequential analyses using electrospray ionization mass spectrometry (ESI–MS). Here we show that capillary electrophoresis (CE) can avoid that tradeoff by organizing predominantly singly charged small molecules and multiply charged peptides into partially resolved, analyte-class-dependent regions of migration time–m/z space. Leveraging this intrinsic electrophoretic organization together with charge- and m/z-resolved precursor selection, we developed a single-run CE–ESI–MS workflow that combines single-vial sample processing with class-resolved tandem MS acquisition. In a HeLa digest spiked with 17 amino acids, the integrated analysis detected all amino acids while preserving proteomic depth relative to a dedicated proteomics run, yielding 1,221 versus 1,227 cumulative protein groups. Applied to identified single *Xenopus laevis* blastomeres, the method provided matched readouts of 86 metabolite features together with 1,097 and 1,083 protein groups from D1.1 and V1.1 cells, respectively. The paired measurements resolved cell-type-dependent molecular differences and mapped protein and metabolite changes into shared pathway context. These results establish analyte-class-dependent electrophoretic organization coupled to class-resolved MS acquisition as an analytical basis for single-run proteome–metabolome analysis by CE–ESI–MS in material-limited samples.

## INTRODUCTION

Biological state emerges from coordinated variation across molecular layers, making same-specimen proteome–metabolome characterization a central analytical objective.^[1–3]^ This challenge is acute for material-limited samples, in which protein-derived peptides and polar small molecules differ in abundance, charge state, chemical diversity, ionization behavior, and sample-handling requirements.^[4–6]^ Existing workflows typically manage this incompatibility by splitting samples, processing analyte classes sequentially, or partitioning material into protein- and metabolite-enriched fractions, but these steps can decouple molecular readouts, increase analyte loss or contamination, and impose class-specific optimization burdens as input decreases.^[6–7]^ Even workflows that recover both classes from a shared starting material must still reconcile extraction chemistry, matrix effects, and downstream compatibility across analyte classes.^[5–7]^

Single-injection MS strategies have begun to address this measurement-level limitation by using class-resolved elution, polarity switching, tailored acquisition windows, hybrid acquisition schemes, gas-phase ion mobility, or computational deconvolution to acquire chemically diverse analytes in one experiment. Examples include LC-based MOST-MS for proteome–lipidome analysis, scPMA for concurrent peptide–metabolite acquisition, and direct-infusion SMAD-MS for extracted protein and moderately polar to apolar metabolites.^[8–10]^ Collectively, these approaches demonstrate the feasibility of single-injection multi-omics, while highlighting a central analytical requirement: chemically diverse analytes must be organized before or during MS detection to limit ionization suppression, precursor competition, spectral congestion, and ambiguous feature assignment.^[11–12]^ Front-end separation provides this organization by distributing analytes and matrix components across time while adding an identifying coordinate that complements accurate mass and tandem MS (MS^2^) information.^[11]^

Capillary electrophoresis (CE) offers a separation principle well matched to this requirement because electrophoretic mobility depends on hydrodynamic radius and charge rather than hydrophobic retention.^[11–12]^ In peptide–metabolite mixtures, this creates an analytical asymmetry: polar small molecules are commonly detected as singly charged, low-m/z ions, whereas tryptic peptides are commonly detected as multiply charged ions distributed across broader migration time (MT)–m/z domains. CE can therefore, in principle, translate intrinsic chemical differences between analyte classes into partially resolved analytical space before MS detection. Although CE–ESI–MS has enabled sensitive, high-efficiency microscale proteomic and metabolomic measurements as separate applications,^[12–14]^ its use for single-run proteome–metabolome analysis has not yet been demonstrated.

Here we tested whether analyte-class-dependent electrophoretic organization can be paired with charge- and m/z-resolved precursor selection to acquire metabolite and peptide tandem MS (MS^2^) spectra from one CE separation. Predominantly singly charged polar small molecules and multiply charged peptides occupied partially resolved domains of MT–m/z space; although not fully separated by migration time alone, their combined differences in MT, m/z, and charge state provided sufficient analytical structure for integrated acquisition. We implemented this strategy through single-vial metabolite and protein processing, followed by class-resolved MS^2^ acquisition in a single CE–ESI–MS run. We first benchmarked class organization, molecular coverage, reproducibility, and quantitative performance against dedicated mono-omics workflows using amino acid and proteome standards. We then demonstrated the approach in material-limited biological specimens by analyzing identified single blastomeres from *Xenopus laevis* embryos and testing whether matched protein and metabolite-feature profiles from the same preparation could resolve cell-type-dependent molecular states.

## RESULTS AND DISCUSSION

As outlined in **Figure 1**, this work tests whether differences in MT, m/z, and charge state can be coupled to class-resolved MS^2^ acquisition so that polar small molecules and peptides can be measured from one CE–ESI–MS separation. Because polar small molecules are detected predominantly as singly charged ions whereas peptides are observed mainly as multiply charged ions, one separation can support acquisition logic for both molecular classes. We first established this concept in defined standards, then applied it to single embryonic-cell preparations to assess performance with biologically complex, input-limited material.

**Figure 1.**
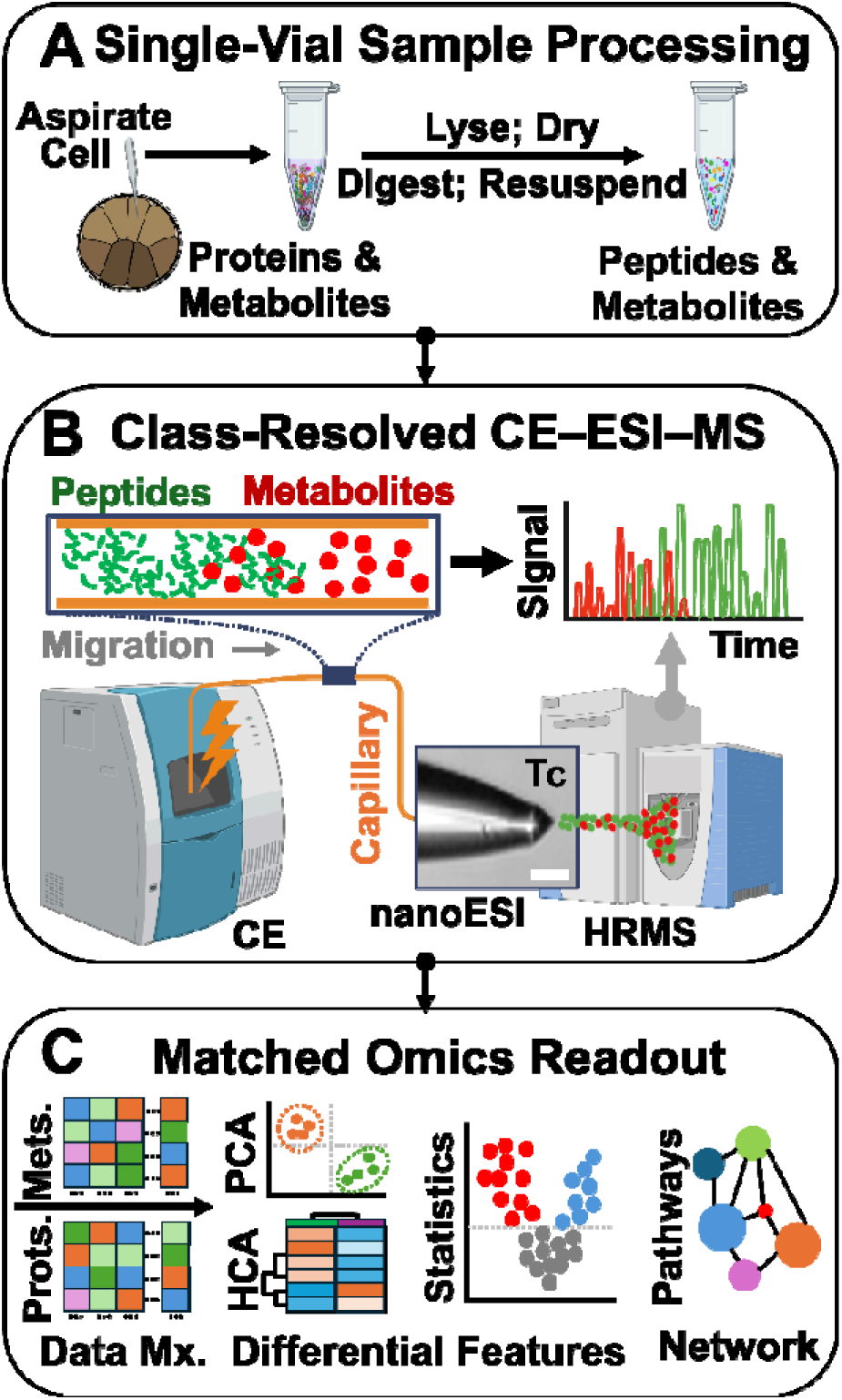
Concept and workflow for single-run proteome–metabolome analysis by CE–ESI–MS. **(A)** A D1.1 or V1.1 blastomere from a 16-cell *X. laevis* embryo was capillary microsampled and processed in one vial to co-recover metabolites and proteins, followed by protein digestion to produce a peptide–metabolite mixture. **(B)** The mixture was analyzed in a single CE–ESI–MS run, in which migration time organization, charge-state filtering, and m/z-resolved MS detection generated class-resolved metabolite and peptide signals. **(C)** PGs (Prots.) and metabolite features (Mets.) were quantified as separate abundance matrices (Mx.) from the same preparation, then analyzed for differential features using chemometric data analysis and pathway-level relationships. Key: HCA, hierarchical cluster analysis; HRMS, high-resolution mass spectrometer; nanoESI, nano-flow electrospray ionization source; PCA, principal component analysis; Tc, Taylor cone.

### Electrophoretic organization produces partially resolved small-molecule and peptide domains

The analytical framework in **Figure 1** required a single CE condition that preserved measurable class-dependent migration behavior while remaining compatible with both polar small molecules and peptides. To test this requirement, we combined 17 amino acids, used as representative polar metabolites, with a HeLa proteome digest and benchmarked single-run dual-omics analysis against a two-run mono-omics reference workflow in which amino acids and peptides were analyzed separately under otherwise matched conditions (**SI Methods**). All measurements were performed by CE–ESI–MS using the same platform and separation conditions. This design isolated the central analytical question: whether representative polar metabolites and peptides retained sufficient analyte-class-dependent organization in MT–m/z space to enable a single integrated acquisition.

To support simultaneous analysis of both classes, the CE conditions had to preserve peptide compatibility while maintaining small-molecule separation. Based on prior work,^[15]^ we used 1 M formic acid with 25% v/v acetonitrile as the background electrolyte to suppress deprotonation of silanol groups on the inner fused-silica capillary wall and minimize peptide adsorption. Samples were prepared in 0.5% v/v acetic acid and 50% v/v acetonitrile to reduce conductivity and promote field-amplified sample stacking. Under these conditions, the 17 amino acids were detected predominantly between about 30–50 min and *m/z* 75–250 Th (**Fig. 2A**, **Table S1**), whereas peptides populated electrophoresis-correlative trend lines^[16–18]^ spanning ∼25–65 min and *m/z* 200–1,500 (**Fig. 2B**, **Table S2**). Despite partial temporal overlap, the amino acids remained concentrated in a low-m/z, predominantly singly charged analytical domain, whereas peptides extended across broader MT and m/z ranges as multiply charged ions. Thus, CE did not fully separate the two classes by migration time alone, but it produced analyte-class-dependent MT–m/z organization that could be combined with charge-state and m/z filtering for class-resolved acquisition.

**Figure 2.**
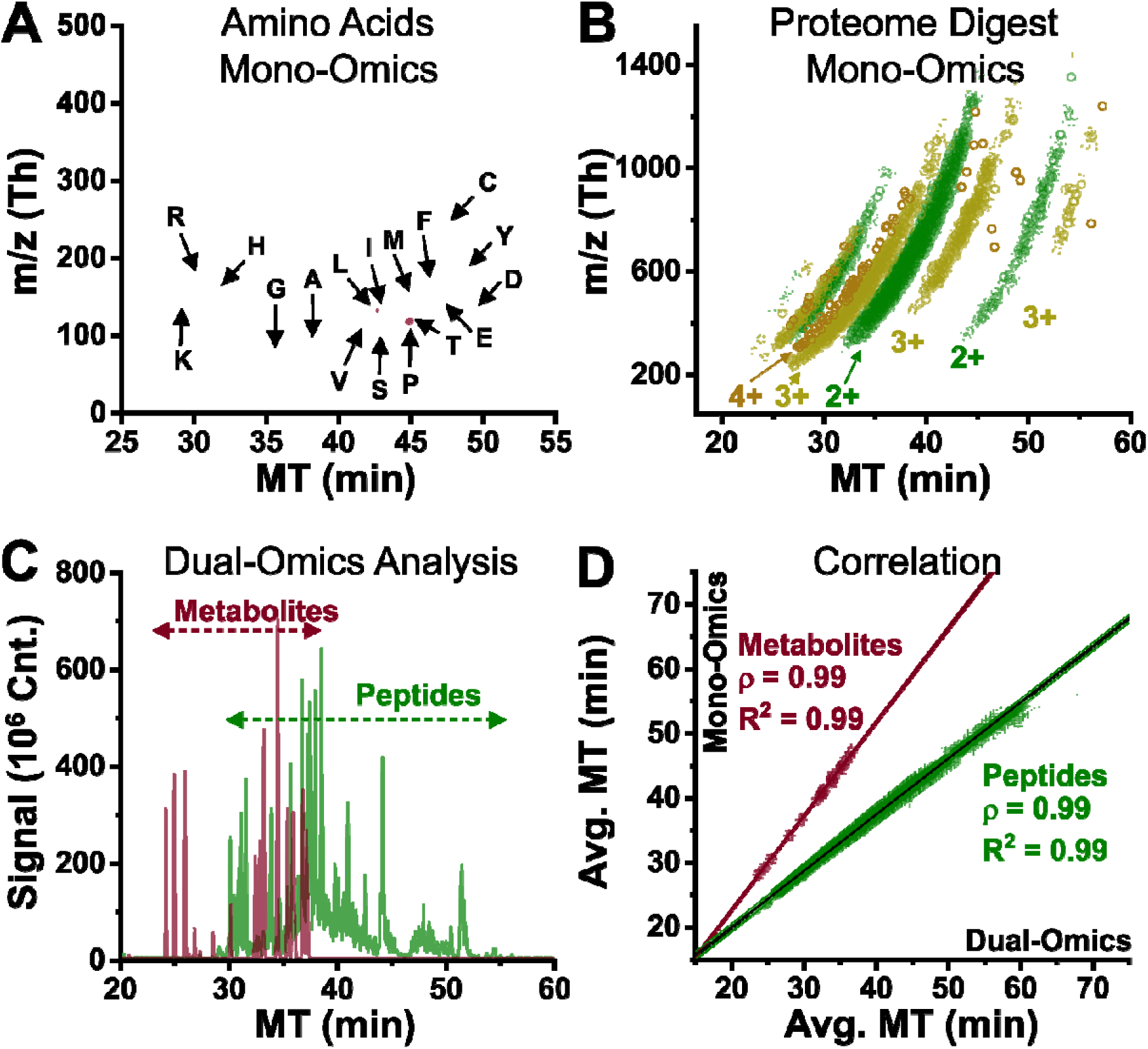
Electrophoretic organization and preservation of class-resolved behavior in single-run CE–ESI–MS. **(A)** Migration time (MT) versus m/z distribution of 17 amino acid standards used as representative polar metabolites, showing a compact low-m/z analytical domain. **(B)** MT versus m/z distribution of peptides from the HeLa proteome digest, which populated broader electrophoresis-correlative trend lines at higher m/z. **(C)** Single-run dual-omics analysis of the mixed amino acid–HeLa digest sample, shown as overlaid extracted-ion electropherograms for the amino acids and the peptide base-peak electropherogram (BPE; *m/z* 500–1,500 Th), demonstrating partial temporal overlap but different dominant signal domains for amino acids and peptides. **(D)** Correlation of average (Avg.) MT values measured in the two-step reference analyses and the single-run dual-omics analysis for amino acids and peptides, showing that co-analysis preserved the separation behavior of both analyte classes. Error bars (RSD) are within symbols. Pearson correlation (ρ) and linear regression (R^2^) coefficients are shown. Shaded bands denote 95% confidence intervals. Amino acids are labeled using standard one-letter abbreviations.

To determine whether this organization was retained during co-analysis, the HeLa digest and amino acid mixture were combined into a single sample and analyzed in one CE–ESI–MS run. **Figure 2C** shows overlaid extracted-ion electropherograms for the amino acids together with the peptide base-peak electropherogram restricted to *m/z* 500–1,500 Th, demonstrating partial temporal overlap but different dominant signal domains for amino acids and peptides. The detected amino acids and peptides are listed in **Tables S1** and **S2**. To assess whether co-analysis altered separation behavior, we compared MT values from the reference analyses with those from the single-run dual-omics analysis (**Fig. 2D**). For both classes, MT values were highly correlated between the two acquisition formats, indicating that co-analysis did not measurably disrupt separation behavior (amino acids, Pearson ρ = 0.99; peptides, ρ = 0.99). Across the technical replicates, the average MT reproducibility was ∼5% CV for the amino acid-only analysis and ∼2% CV for the dual-omics analysis. These results show that mixed analysis maintained the analyte-class-dependent organization and high reproducibility required for class-resolved acquisition from a single integrated CE–ESI–MS run.

### Class-resolved MS^2^ preserves proteomic depth and quantitative performance in a single CE–ESI–MS run

The class-dependent organization in **Figure 2** provided the basis for acquisition-level class resolution within a single CE–ESI–MS analysis. Because MT alone did not fully separate small molecules and peptides, the MS^2^ acquisition method used charge state and m/z range to distinguish precursor classes. The method alternated between two MS^1^scans: a small-molecule survey that selected singly charged precursors over *m/z* 50–500 Th and a peptide survey that selected multiply charged precursors over *m/z* 200–1,500 Th. Precursors from each survey were then subjected to data-dependent MS^2^ under matched fragmentation conditions (**SI Methods**). Thus, charge state and m/z range served as acquisition logic that translated electrophoretic organization into class-resolved MS^2^.

This strategy preserved proteomic identification depth relative to the reference workflow. Across technical triplicates, protein group (PG) identification depth was statistically comparable between the protein-only reference workflow (887 PGs per run) and the single-run dual-omics workflow (892 PGs per run; raw *p* = 0.66, Mann–Whitney U test; **Fig. 3A**). Cumulative identifications were similar, with 1,227 PGs in the reference workflow and 1,221 PGs in the single-run dual-omics workflow. Both workflows detected all 17 amino acids. The workflows identified 977 PGs in common, corresponding to 66.4% of the combined protein identifications, with similar fractions unique to the reference and dual-omics workflows (**Fig. 3B**). Thus, class-resolved acquisition enabled amino acid and peptide MS^2^ readout in one CE–ESI–MS run while maintaining proteomic identification depth.

**Figure 3.**
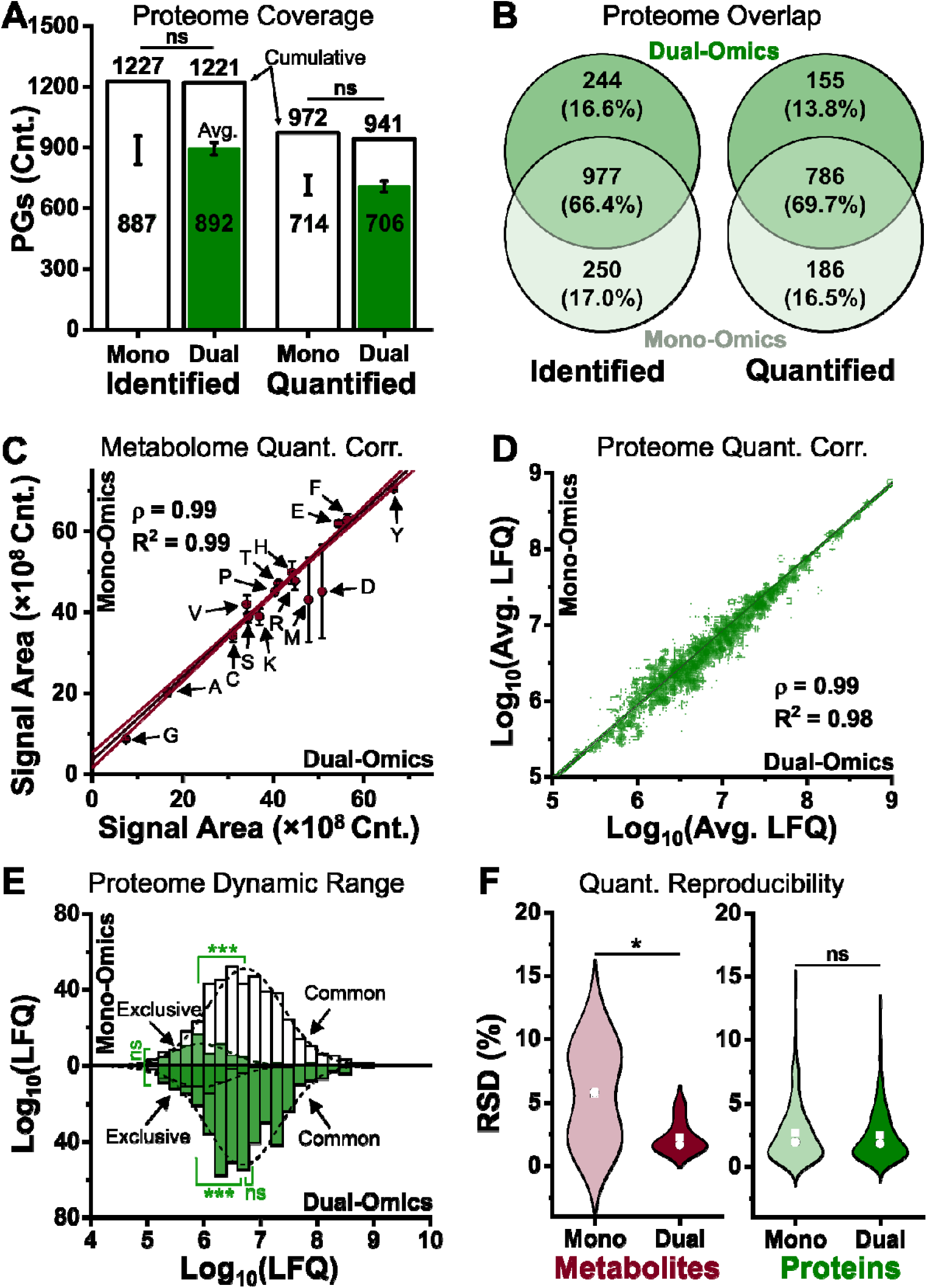
Single-run dual-omics analysis preserves proteomic depth and quantitative performance in a representative amino acid–peptide mixture. **(A)** Per-run and cumulative PG identification and label-free quantification (LFQ) depth for the protein-only reference and single-run dual-omics workflows. **(B)** Overlap of identified or quantified PGs between workflows. Correlation (Corr.) between **(C)** metabolite median-normalized signal areas and **(D)** protein LFQ abundance values. **(E)** LFQ abundance distributions of PGs detected exclusively or commonly between workflows. **(F)** Precision of amino acid and protein quantification, expressed as relative standard deviation (RSD, %). Statistics were calculated using the Mann–Whitney U test: ns, not significant (*p* ≥ 0.05); *, raw *p* < 0.05; ***, raw *p* < 0.001; circle, mean; square, mean. Key: ρ, Pearson correlation coefficient; R², coefficient of linear regression. Confidence bands show 95%. Amino acids are labeled using standard one-letter abbreviations.

Identification depth and quantitative coverage report distinct aspects of method performance; we therefore compared the number of reproducibly quantified features between the dual-omics and mono-omics workflows. For proteins, LFQ produced statistically comparable quantification depths between workflows, averaging 714 quantified PGs per run in the reference workflow and 706 PGs per run in the single-run dual-omics workflow (raw *p* = 0.99, Mann–Whitney U test; **Fig. 3A**). Cumulative LFQ coverage was similar, with 972 and 941 PGs quantified in at least 1 of 3 TRs for reference and single-run analyses, respectively (**Fig. 3A**, **Table S3**). Among cumulatively quantified PGs, 69.7% were shared between workflows, with similar workflow-specific fractions (**Fig. 3B**, **Table S3**). Missing-value rates did not differ significantly between the workflows (raw *p* = 0.38, Mann–Whitney U test; **Fig. S1**). Thus, expanding the acquisition method to include small-molecule precursors did not measurably compromise the completeness of the LFQ dataset.

Comparable quantification depth alone does not establish quantitative equivalence; the measured abundances must also agree across workflows while preserving dynamic-range coverage and precision. Amino acid signal areas were strongly correlated between the reference and single-run analyses, with Pearson’s ρ = 0.99 and R² = 0.99 (**Fig. 3C**). Among PGs quantified in both workflows, LFQ abundances were likewise strongly correlated (Pearson’s ρ = 0.99, R² = 0.98; **Fig. 3D**). The abundance distributions of PGs detected exclusively or commonly between workflows were statistically comparable, supporting similar dynamic-range coverage (**Fig. 3E**). Quantitative precision was likewise retained, with mean and median relative standard deviations below 10% for both amino acids and PG LFQs (**Fig. 3F**). PG LFQ precision remained statistically comparable between workflows (raw *p* = 0.53, Mann–Whitney U test), whereas amino acid measurements exhibited significantly lower relative standard deviations in the dual-omics workflow than in the mono-omics reference (raw *p* = 0.011, Mann–Whitney U test). Thus, class-resolved CE–ESI–MS expanded molecular readout while maintaining proteomic depth, quantitative coverage, and measurement precision.

### Single-cell CE–ESI–MS dual-omics yields paired molecular coverage from lineage-validated *X. laevis* blastomeres

The analytical performance established with standards provided the basis for applying the workflow to biologically complex, input-limited preparations. We selected the left dorsal-animal midline (D1.1) and left ventral-animal midline (V1.1) blastomeres from 16-cell *X. laevis* embryos as test specimens (**Fig. 4A**). These cells are reproducibly identifiable by position and pigmentation, preferentially give rise to neural and epidermal lineages, respectively,^[19]^ and have shown metabolomic^[20]^ and proteomic^[15]^ differences in previous single-omic studies. To validate prospective cell assignment, D1.1 and V1.1 blastomeres identified by morphological criteria were injected with fluorescent dextran, and descendant fluorescence patterns were examined at the larval stage (**Fig. 4A**). Intracellular contents were microaspirated from individual D1.1 and V1.1 blastomeres, each from separate live embryos, then quenched in cold methanol and processed in a single vial to minimize analyte loss (**Fig. 4B**; **SI Methods**). For each cell type, five biological replicates were analyzed, with each biological replicate prepared from one embryo and measured by three technical CE–ESI–MS injections. The same vial was carried through protein digestion and reconstitution, and each injection consumed ∼0.25% of a biological replicate.

**Figure 4.**
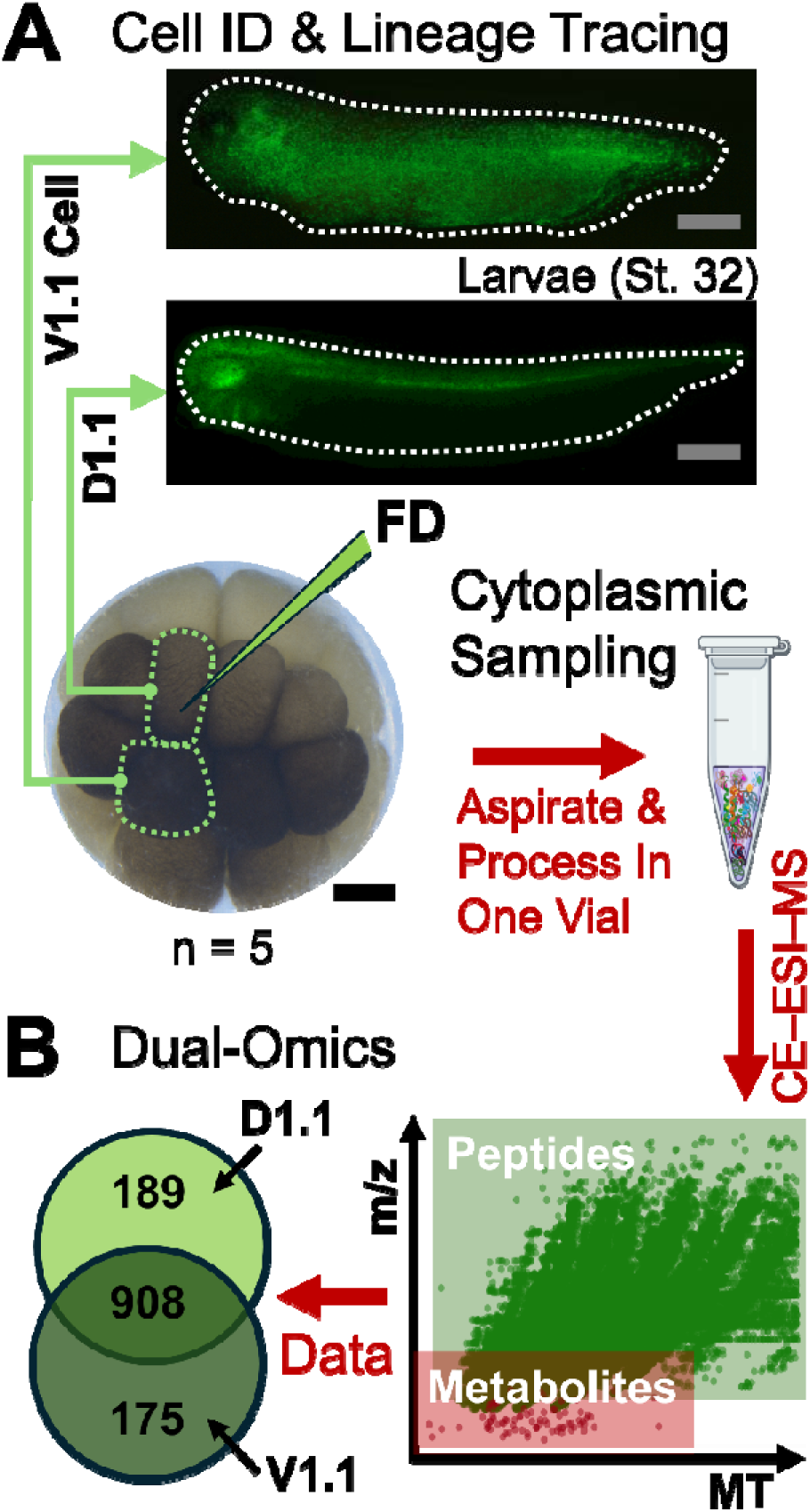
Cell-lineage validation enables targeted dual-omics CE–ESI–MS of identified embryonic blastomeres. **(A)** D1.1 and V1.1 blastomeres in the 16-cell embryo were prospectively identified using established morphological criteria, and their identities were validated by fluorescent dextran (FD) lineage tracing to descendant cells at the larval stage. **(B)** Validated cellular material was aspirated and processed in a single vial for single-run dual-omics CE–ESI–MS, linking defined embryonic cell identity to peptide and metabolite detection from the same preparation. Peptide and metabolite features detected were visualized in m/z–MT space to show the molecular coverage obtained from the same CE–ESI–MS analysis (**right**). The characterized D1.1 and V1.1 proteomes were compared by overlap analysis (**left**). Scale bars: black, 250 µm; gray, 400 µm.

The single-run workflow yielded paired metabolic and proteomic readouts from the same lineage-validated blastomere preparations. Peptide and metabolite features occupied partially resolved regions of m/z–MT space (**Fig. 4B**), illustrating the molecular coverage obtained from the same CE–ESI–MS analysis. In total, 86 metabolite features were matched to a prior single-blastomere reference set by accurate m/z and MT with nonlinear alignment (**Table S4; SI Methods**). In the same analyses, 1,097 and 1,083 PGs were identified from D1.1 and V1.1 blastomere preparations, respectively (**Table S5**). This molecular coverage is consistent with prior single-cell metabolic^[20–22]^ and proteomic^[15^, ^22–24]^ measurements on these blastomere phenotypes. Across cell types, all 86 metabolite features were detected in both D1.1 and V1.1 preparations, whereas overlap analysis showed that most PGs were shared between D1.1 and V1.1 preparations, with smaller subsets detected only in one preparation **(Fig. 4B**). Protein abundances spanned ∼6 orders of magnitude (**Table S6**), and metabolite MF abundances spanned ∼3 orders of magnitude, further supporting broad molecular coverage from the input-limited preparations **(Fig. S2**). These coverage metrics supported quantitative profiling of the lineage-validated cell types.

### Matched proteome–metabolome abundances distinguish D1.1 and V1.1 molecular states

Because D1.1 and V1.1 identities were assigned morphologically before sampling, the critical analytical test was whether paired abundance profiles retained sufficient structure to distinguish these blastomere states. Separate hierarchical cluster analysis of metabolite-feature peak areas and PG LFQ abundances from the single-run dual-omics workflow organized samples by cell type and revealed D1.1- and V1.1-enriched abundance patterns (**Fig. 5A,B**). Differential analysis provided a complementary global view of these cell-type-dependent abundance patterns. Overall, 22 metabolite features and 82 PGs differed significantly between D1.1 and V1.1 blastomeres at raw *p* < 0.05, whereas most quantified measurements remained comparable between cell types (**Fig. 5A,B**; **Table S7**).

**Figure 5.**
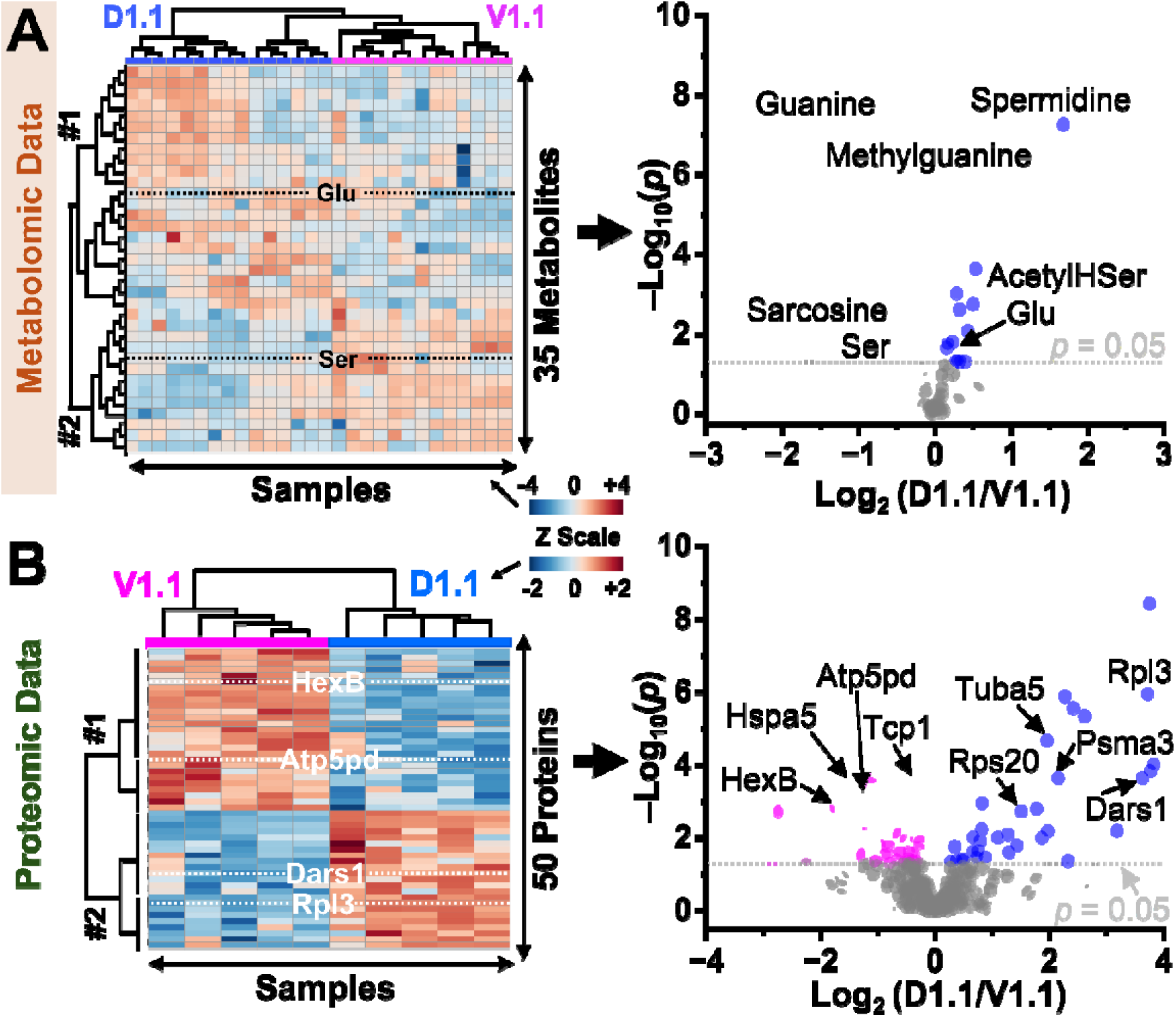
Matched metabolite and proteomic abundance profiles distinguish D1.1 and V1.1 blastomere molecular states from single-run CE–ESI–MS dual-omics measurements. **(A)** Hierarchical cluster analysis (HCA) of selected metabolite features (left) and volcano plot (right) of all quantified metabolite features. **(B)** HCA–heatmap of selected PGs (left) and volcano plot of all quantified PGs (right). In the volcano plots, blue points mark D1.1-enriched features or PGs, magenta points mark V1.1-enriched features or PGs, and gray points mark nonsignificant measurements; dashed horizontal lines mark raw *p* = 0.05 (Mann–Whitney U test). Key: AcetylHSer, acetylhomoserine; Glu, glutamic acid; LFQ, label-free quantification; Ser, serine.

Several abundance differences were consistent with previously reported molecular distinctions between these blastomere states. Serine signal abundance and the mitochondrial protein Atp5pd were elevated in V1.1 blastomeres, consistent with earlier single-omics measurements of these cell types (**Fig. 5**; **Table S7)**.^[15^, ^17, 22]^ Among lineage-enriched detections, Tubb4a was observed only in D1.1 blastomeres, consistent with a neural-lineage association^[25]^, whereas Ptbp1, a repressor of neuronal mRNAs,^[26–27]^ was observed only in V1.1 blastomeres, consistent with prior reports in *Xenopus* epidermal cells.^[28]^ These results show that class-resolved CE–ESI–MS dual-omics can recover paired metabolite-feature and proteomic abundance signatures from individual embryonic cells and use them to resolve cell-type-dependent molecular states.

Pathway-level analyses further placed the paired measurements in biochemical context. STRING analysis of all identified PGs provided a protein-centered view of the interaction networks and functional modules captured by the single-cell CE–ESI–MS workflow, including proteostasis, ribosome, oxidative phosphorylation, and carbon metabolism modules (**Fig. 6A**). Joint pathway analysis of differential PGs and metabolite features mapped the cell-type-dependent changes onto 61 biochemical pathways after removal of disease- and virus-associated entries (**Table S8**). These pathways included arginine–proline metabolism, glutathione metabolism, glycine–serine–threonine metabolism, glycolysis/gluconeogenesis, one-carbon pool by folate, pentose phosphate pathway, and the TCA cycle (**Fig. 6B**; **Table S8**). Thus, the same single-run CE–ESI–MS measurements supported both broad proteomic pathway coverage and dual-layer mapping of protein and metabolite differences.

**Figure 6.**
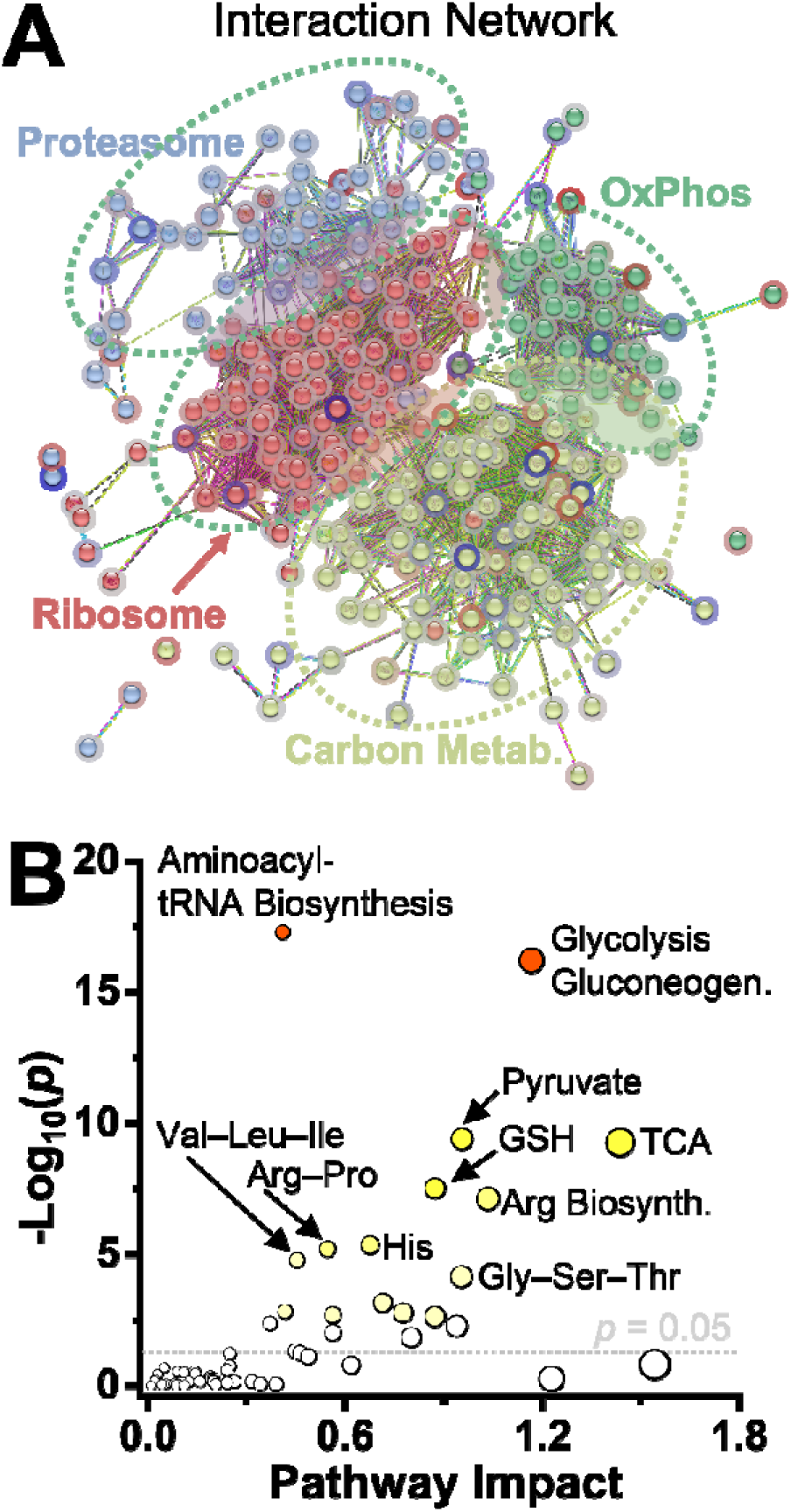
Pathway-level analyses show broad proteomic coverage and dual-layer biochemical mapping from D1.1 and V1.1 blastomeres. **(A)** STRING protein-protein interaction analysis of all identified PGs showed broad functional coverage across major protein networks, including proteostasis, ribosome, oxidative phosphorylation (OxPhos), and carbon metabolism. **(B)** MetaboAnalyst joint pathway analysis mapped differential PGs and metabolite features onto 61 biochemical pathways after removal of disease- and virus-associated entries. Representative pathways are labeled using three-letter amino acid abbreviations. Dashed horizontal line marks raw *p* = 0.05 (Mann–Whiney U test). Key: Biosynth., biosynthesis; GSH, glutathione; TCA, tricarboxylic acid cycle. Amino acids are labeled using standard three-letter abbreviations.

The serine/one-carbon metabolic neighborhood provided a representative example of this cross-layer interpretation. Serine signal abundance was elevated in V1.1 blastomeres at the metabolite-feature level, whereas acetylhomoserine and formimidoyltransferase cyclodeaminase (FTCD), an enzyme linking histidine catabolism to folate-dependent one-carbon metabolism, were higher in D1.1 blastomeres. Additional mapped features connected this neighborhood to nucleotide biosynthesis, including bifunctional purine biosynthesis protein ATIC, which uses a folate-derived one-carbon unit in *de novo* purine biosynthesis, together with glycine, methionine, and glycine betaine (**Fig. 6B**; **Table S7**). These pathway relationships should not be interpreted as direct measurements of metabolic flux or developmental mechanism. Rather, they demonstrate that class-resolved CE–ESI–MS links metabolite-feature abundance and PG abundance from the same cellular extracts, extending single-cell CE–ESI–MS from parallel molecular detection to integrated biochemical interpretation in early *X. laevis* embryos.

## CONCLUSIONS

These results show that CE need not serve only as a front-end separation for one molecular class at a time. By coupling analyte-class-dependent electrophoretic organization to charge-and m/z-resolved MS acquisition, a single electrophoretic separation supported parallel readout of chemically distinct analytes without an evident penalty to proteomic performance. In standards, this strategy preserved amino acid detection, proteomic identification depth, LFQ coverage, and quantitative agreement relative to dedicated mono-omics measurements. In identified single *X. laevis* blastomeres, the same workflow provided matched metabolite and protein-group readouts from the same preparation and enabled pathway-level integration of both molecular classes.

This strategy addresses a central limitation in input-limited proteome–metabolome analysis by preserving matched molecular information while avoiding sample splitting or sequential measurements. The current implementation provides metabolite feature coverage rather than comprehensive metabolite annotation, and the pathway results should be interpreted as biochemical context rather than direct flux measurements. Future extensions should focus on improving metabolite identification confidence, expanding small-molecule and proteome coverage through CE-compatible acquisition strategies, and automating low-input sample processing to increase throughput and reproducibility. These results establish analyte-class-dependent electrophoretic organization coupled to class-resolved MS acquisition as a practical route to matched proteome–metabolome readout from scarce biological samples in a single CE–ESI–MS analysis.

## Acknowledgement

Parts of this research were supported by the Chan Zuckerberg Initiative (award no. DAF2023-331951 to P.N.) or the National Institute of General Medical Sciences of the National Institutes of Health (award no. R35GM124755 to P.N.).

## Author Contributions

P.N. conceptualized and supervised the study. R.M.O. cultured the embryos and performed single-cell microsampling. R.K. developed the analytical methodology, processed the samples, and acquired the MS data. R.K. and P.N. analyzed and interpreted the proteomics data. R.M.O. and P.N. analyzed and interpreted the metabolomics data. R.K. drafted the manuscript with assistance from R.M.O. P.N. edited and finalized the manuscript. P.N. secured the funding. All authors reviewed and approved the final manuscript.

