## Supplementary material for "Analyte-Class-Dependent Electrophoretic Organization Enables Single-Run Proteome–Metabolome Analysis by CE–ESI–MS": SI Document

### SI MATERIALS AND METHODS

**Materials.** Acetic acid (AcOH), acetonitrile (ACN), formic acid (FA), and methanol (MeOH) were LC–MS grade and purchased from Fisher Scientific (Hampton, NH). Ammonium bicarbonate was obtained from Avantor (Center Valley, PA). The amino acid standard mixture (AAS18) was purchased from Millipore Sigma (Burlington, MA) and contained L-alanine, ammonium chloride, L-arginine, L-aspartic acid, L-glutamic acid, glycine, L-histidine, L-isoleucine, L-leucine, L-lysine, L-methionine, L-phenylalanine, L-proline, L-serine, L-threonine, L-tyrosine, and L-valine at 2.5 mM each, and L-cystine at 1.25 mM. The HeLa proteome digest (part no. 88329) was obtained from Thermo Fisher Scientific (Waltham, MA). Fluorescent dextran (FD; dextran-conjugated Alexa Fluor 488, Mw = 3,000 g/mol, part no. D34632) was purchased from Invitrogen (Carlsbad, CA). All other reagents were purchased from Sigma-Aldrich (St. Louis, MO) unless otherwise noted. For method development, 5 µg of HeLa digest was dissolved in 10 µL of the amino acid mixture to yield final amino acid concentrations of 10 µM (L-cystine, 5 µM) in sample solvent.

**Parts.** Capillary electrophoresis (CE) separations were performed using bare fused-silica capillaries (40/105 µm inner/outer diameter; part no. 1068150596) from Polymicro Technologies (Phoenix, AZ) without further modification. CE–electrospray ionization (CE–ESI) sources were fabricated from borosilicate glass capillaries (0.75/1.00 mm; part no. B100-75-10) from Sutter Instrument (Novato, CA).

**Solutions.** For embryo culture, 100% Steinberg's solution was prepared as described elsewhere.[1] Lineage tracing solution was 0.5% v/v FD in DEPC-treated water. CE–ESI–MS analyses used the following solvents: *sample solvent*, 0.5% v/v AcOH in 50% v/v ACN in water; *background electrolyte* (*BGE*), 1 M FA in 25% v/v ACN in water; and CE–ESI *sheath solution*, 0.5% v/v AcOH in 10% v/v ACN in water.

**Animal Care.** Sexually mature adult *Xenopus laevis* frogs were obtained from Xenopus1 (Dexter, MI). All protocols ensuring the humane care and use of the animals were approved by the Institutional Animal Care and Use Committee at the University of Maryland, College Park (approval no. R-FEB-24-05). Embryos were produced by gonadotropin-induced natural mating of two parental pairs following established protocols.[1]

**Cell Identification and Lineage Tracing**. Two-cell stage embryos with stereotypical pigmentation were selected and cultured to the 16-cell stage to enable accurate identification of the dorsal-animal midline (D1.1) and ventral-animal midline (V1.1) cell types based on pigmentation, size, and location in reference to established cell-fate maps.[2] For lineage tracing, D1.1 and V1.1 cells were injected with FD and cultured following previously published research.[3]

**Single-Cell Sampling and Processing.** The contents of D1.1 and V1.1 blastomeres were collected from live embryos in randomized order by capillary microsampling following established protocols.[4-7] The subcellular aspirates were discharged into individual LoBind Eppendorf (Enfield, CT) microvials containing 10 µL of MeOH, then stored at –80 ºC. For analysis, the cellular extracts were thawed, vacuum-dried, and resuspended in 4 µL of 50 mM ammonium bicarbonate, followed by rapid heat denaturation (60 °C for 20 min) before digestion (1 µg MS-grade trypsin, 37 °C, 5 h) without reduction, alkylation, or desalting, as established.[6]

The resulting peptide–metabolite mixtures were vacuum-dried for storage at –80 °C. The mixtures were thawed and reconstituted in 4 µL of CE–ESI–MS sample solvent prior to analysis.

**CE–ESI–MS Analysis**. ***CE–ESI.*** Separations were performed on a commercial CE platform (CESI 8000 Plus, AB SCIEX, Framingham, MA) using an electrokinetically pumped sheath-flow CE–ES interface[8] operated in the cone-jet regime[9] for maximal ionization efficiency[10]. Experimental parameters were as follows: capillary length, 145 cm; separation potential, +25 kV applied at the inlet end of the BGE-filled CE capillary; emitter tip diameter, 30 µm; CE–electrospray regime, cone-jet (confirmed based on microscopy and total ion current analysis[10]). Between runs, the capillary was conditioned with 0.1 M NaOH for 10 min, rinsed with water for 5 min, and equilibrated with BGE for 10 min. Blank injections of BGE yielded no detectable protein identifications, indicating negligible carryover. ***MS***. Ions were detected using an Orbitrap mass spectrometer (Q Exactive Plus, Thermo Fisher Scientific, Waltham, MA). Mass (m/z) accuracy was ensured by calibrating the instrument prior to data acquisition according to manufacturer-recommended procedures. Data-dependent acquisition was used to alternate $MS^1$ survey scans and tandem MS ($MS^2$) scans for peptide- and metabolite-like signals. Metabolite scans targeted singly charged species over *m/z* 50–500 Th, excluding unassigned and multiply charged ions. Peptide scans targeted multiply charged species over *m/z* 200–1,500 Th, excluding unassigned, singly charged, and >+8 ions. This charge-state filtering confined metabolites and peptides to separate acquisition cycles and minimized cross-selection of precursors between analyte classes. $MS^1$ spectra were acquired at 70,000 FWHM resolution with an AGC target of $1 \times 10^6$ counts and a maximum injection time of 50 ms. Selected precursors from both survey scans were subjected to $MS^2$ fragmentation. $MS^2$ spectra were acquired over *m/z* 200–2,000 Th at 17,500 FWHM resolution, with an AGC target of $5 \times 10^4$ counts, maximum injection time of 100 ms, and an isolation window of 1.5 Th. Fragmentation was performed using higher-energy collisional dissociation at 28% NCE in nitrogen. All other $MS^2$ acquisition parameters were defined to the default by the instrument control software.

**Data Analysis. *Proteomics.*** MS data were analyzed using Proteome Discoverer 3.0 (Thermo Scientific) with SEQUEST HT against the human HeLa proteome (UniProt UP000005640, 20,309 entries, downloaded June 8, 2021) or the *X. laevis* proteome (UniProt UP000186698, 33,996 entries, downloaded April 4, 2023). Search parameters were: minimum peptide length, 6 aa; maximum missed cleavages, 2; precursor mass tolerance, 10 ppm; fragment mass tolerance, 20 mTh; static modification, Cys carbamidomethylation (HeLa) or none (*X. laevis*); dynamic modification, methionine oxidation; all other parameters, default. Peptide and protein identifications were filtered to <1% false discovery rate using a reversed-sequence decoy database. Proteins annotated in the Common Repository of Adventitious Proteins (cRAP)[11] were removed and are not reported (www.thegpm.org/crap). Protein abundances were estimated by label-free quantification (LFQ) using the Minora algorithm[12] in Proteome Discoverer. HeLa and *X. laevis* proteome LFQ data were median normalized and $\log_{10}$-transformed in MetaboAnalyst 6.0[13]. Proteins were considered quantified when observed in 3 technical replicates (TRs).

***Metabolomics.*** The MS data were processed using Xcalibur 4.1.31.9 (Thermo) with the following parameters: m/z range, *m/z* 50–500 Th; mass tolerance, 50 mDa. A previously established set of *X. laevis* molecular features[4] were extracted using Skyline-daily 25.1.1.258.[14] Skyline $MS^1$ filtering parameters were the following: isotope peaks included, count; precursor mass analyzer, centroided; peaks, 1; mass accuracy, 10 ppm. Skyline instrument parameters were: min m/z, *m/z* 50 Th; max m/z, *m/z* 350 Th; match tolerance, 5 mTh; min time, 10 min; max time, 80 min; precursor adducts, protonated. Name assignment of molecular features were determined utilizing both accurate mass and alignment of migration times using a nonlinear multiparameter regression as described elsewhere[15] and the list of molecular features was from previously published data from our research group. Endogenous metabolites lysine, histidine, valine, threonine, and aspartate were chosen to provide time markers for the nonlinear time regression. Metabolite area data from m/z-selected $MS^1$ scans were median normalized for the standards and sum-normalized, $\log_{10}$-transformed, and auto-scaled for *X. laevis* using MetaboAnalyst 6.0.

**Study Design.** ***Replicates***. For method development and validation, the HeLa proteome digest spiked with 17 amino acids (AAs) was analyzed in three TRs. For biological measurements, n = 5 D1.1 and n = 5 V1.1 *X. laevis* blastomeres (biological replicates, BRs) were analyzed, each in technical triplicate. Each BR was derived from a different embryo from two parental pairs to account for biological variability. Each BR and TR was assigned a unique identifier: $BR_{TR}$.

**Chemometrics**. ***Statistics.*** All statistical analyses were performed in OriginPro 2026 (OriginLab Corp., Northampton, MA). Data normality was assessed using the Shapiro–Wilk test. Normally distributed data were evaluated using Student's t-test, whereas non-normally distributed data were evaluated using the Mann–Whitney U test. Statistical significance was defined as $p < 0.05$. Multiple hypothesis testing correction was performed using the Benjamini–Hochberg method. ***Multivariate Data Analysis.*** Principal component analysis (PCA) was performed using MetaboAnalyst 6.0. ***Outlier Identification (PCA)***. Unsupervised principal component (PC) analysis showed that most TRs clustered tightly. Two samples ($7_1$ and $10_2$; Table S4) were identified as extreme outliers based on large separation from the main cluster (data not shown), with total explained variance of 58.7% (PC1, 36.9%; PC2, 21.8%). Both samples fell outside the 95% confidence ellipses for the D1.1 and V1.1 groups. Because PCA distance measures sample dissimilarity, these deviations suggested either technical artifacts or sample degradation. These samples were excluded from downstream metabolomics analyses.

***Pathway Analysis***. For joint pathway analysis, protein and metabolite fold changes were exported from MetaboAnalyst 6.0. Human orthologs of *X. laevis* genes were assigned using DIOPT[16]. Pathway analysis parameters were: enrichment method, hypergeometric test; topology metric, degree centrality; integration method, combined *p*-values. Protein-protein interaction analysis was performed using STRING[17] with the following settings: database, *X. laevis*; disconnected nodes, hidden; clustering method, k-means (4 clusters).

**Scientific Rigor.** ***Biological Replicates.*** Samples were coded using unique identifiers ($BR_{TR}$) and analyzed in randomized order. Sex as a biological variable was not considered because *X. laevis* embryos do not exhibit sex-specific features at cleavage stages. All biological samples were collected on the same day and analyzed in randomized order. Multivariate analyses were performed blinded to sample identity, which was revealed only after analysis for interpretation. ***Data Filtering.*** For proteomics, proteins were considered quantified within a BR if non-zero LFQ values were present in at least 2 TRs. Across samples, only proteins common to both cell types and quantified in at least 3 out of 5 BRs were retained for downstream analysis. The missing values were imputed using k-nearest neighbors algorithm in MetaboAnalyst 6.0. For cumulative LFQ coverage, proteins quantified in at least 1 of 3 TRs were counted. For RSD/CV precision analysis, only proteins quantified in all 3 TRs were used. Missing value imputation was not applied to preserve analytical rigor for coefficient of variation (CV) calculations. For integrated analyses, only proteins and metabolites detected in both cell types and quantified in at least 3 of 5 BRs were included.

**Safety Considerations.** Standard safety protocols were followed for handling chemicals and biological materials. Personal protective equipment, including gloves and eye protection, was used to mitigate puncture hazards from fused silica capillaries and metal needles. Electrically conductive components of the CE–ESI–MS setup were properly grounded or shielded to prevent electric shock.

### SI TABLES

The following tables are provided as Excel file in the electronic SI document:

**Table S1**. Analysis of amino acid standards via mono-omics and dual-omics.

**Table S2**. HeLa peptides detected using mono-omics and dual-omics.

**Table S3.** HeLa protein identifications.

**Table S4.** Single-cell metabolite features.

**Table S5.** Single-cell protein identifications.

**Table S6.** Single-cell protein quantification.

**Table S7.** Single-cell dual proteome-metabolome profiling.

**Table S8.** Joint proteome-metabolome pathway analysis**.**

#### SI FIGURES

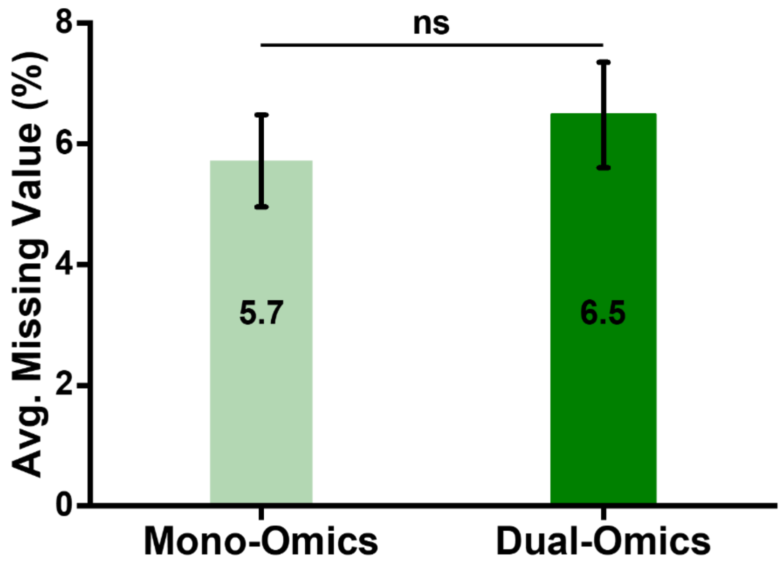

**Figure S1**. Missing-value rate for HeLa proteome digest quantification in mono-omics and dual-omics workflows. The percentage of missing LFQ values was statistically comparable between workflows. Statistics were calculated using the Mann–Whitney U test: ns, not significant (raw $p = 0.38$).

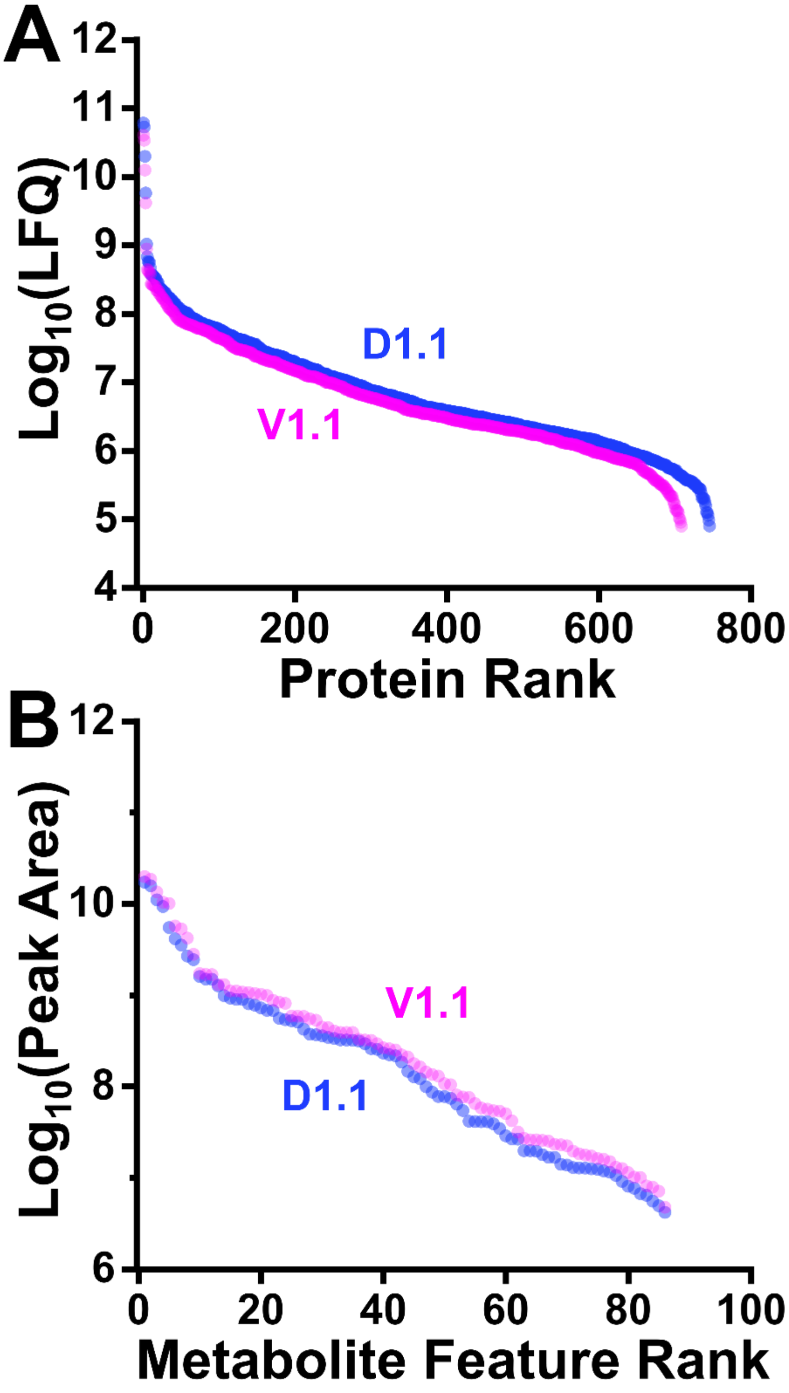

**Figure S2**. Dynamic range of quantification in D1.1 and V1.1 blastomeres by single-run dual-omics CE–ESI–MS. **(A)** Ranked LFQ abundance distributions for PGs. (**B**) Ranked metabolite-feature peak-area distributions.

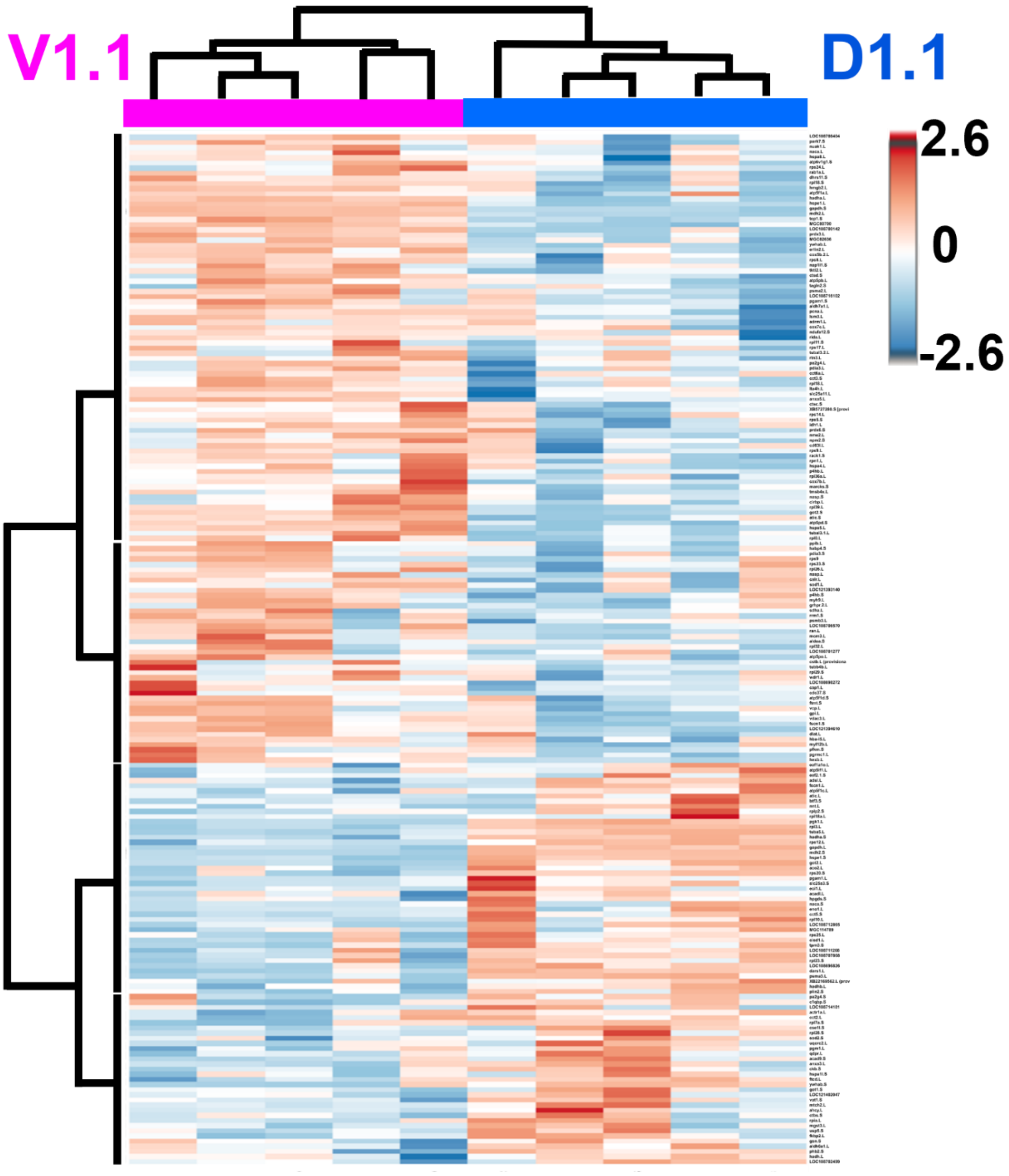

**Figure S3**. Extended hierarchical cluster analysis (HCA) of proteins in D1.1 and V1.1 cells. The top 200 statistically significant proteins clustered samples into distinct cell-type-dependent expression groups.
